# Thermal stimulation of pressure phosphenes

**DOI:** 10.1101/2021.03.12.435166

**Authors:** Alexander Kholmanskiy, Elena Konyukhova, Andrey Minakhin

## Abstract

To investigate effect on the intensity of pressure phosphenes (PP) of various methods of heating hands, as well as manual influence on cervical spine. This study included subjective assessments of the PP intensity in 10 healthy men, and chronometry of bioelectrical activity of brain and heart using electroencephalography (EEG) and electrocardiography (ECG). EEG and ECG frequency spectra respond synchronously to pressure, heat and light. The time of excitation of potential oscillations in the visual area of the cortex of both hemispheres is consistent with the delay in the onset of PP flaring. The effect of PP stimulation is enhanced when the hands and body are heated in a humid environment and at temperatures above 42 °C. Stimulation of PP by heating indicates the convergence of impulses from neurons of the lateral geniculate body (LGB) and nuclei of the thalamus, which are responsible for thermoreceptors in the skin of the palms and fingers. The thermal stimulation effect of PP is enhanced at temperatures above 42 °C due to the fact that thermoreceptors work as pain receptors. The mechanism of PP generation is dominated by the processes of redistribution and recombination of charges in the layers of the retina and LGB.

## 1. Introduction

The development of sensing in living organisms began with mechanoreception, then the organs for the perception of electromagnetic radiation from the Sun and heat appeared. The electromagnetic nature of bioenergy in the process of evolution provided vision to a dominant role in the hierarchy of the nervous system of mammals and humans [1]. This is confirmed by synesthesia of human vision with most of the body’s receptor systems, the total number of which reaches 70 [2, 3]. Table 1 shows the most common types of synesthesia of vision and the percentage of people with them from the total number of surveyed synesthetes (1143 people) [3]. The visual nervous system (VNS) plays a major role in the subordination of homeostasis to the circadian rhythm [4, 5], driven by special light-sensitive retinal ganglion cells and neurons of the suprachiasmatic nuclei [4, 6, 7].

**Table 1.** The percentage of synesthetes with a specific type (1143 synesthetes in total, Data from [3])

| Type of synesthesia | % |
| --- | --- |
| graphemes -> vision | 61.26 * |
| time units -> vision | 22.96 * |
| musical sounds -> vision | 18.05 * |
| general sounds -> vision | 16.21 |
| musical notes -> vision | 7.80 * |
| personalities -> vision | 6.49 * |
| phonemes -> vision | 7.54 * |
| odors -> vision | 6.13 |
| flavors -> vision | 5.78 * |
| pain -> vision | 5.43 |
| touch -> vision | 3.94 |
| emotions -> vision | 3.24 * |
| orgasm -> vision | 1.93 |
| temperatures -> vision | 1.84 |
| lexemes -> vision | 0.70 * |
| kinetics -> vision | 0.53 |
| proprioception -> vision | 0.09 |
| vision -> flavors | 2.98 * |
| vision -> odors | 1.14 |
| vision -> sounds | 3.07 * |
| vision -> touch | 1.58 |

Pressure phosphenes (PP) have been known since the 5th century. BC, in the Middle Ages, hesychasts called phosphenes “Uncreated light”. Electrophosphenes (EPs) [8, 9] were discovered simultaneously with electricity. Typical PPs can be triggered by applying pressure to the eyes with your fingers or by listening to loud sounds after adjusting vision to darkness [10]. It has been established in animals and humans [11, 12, 13, 14] that action potentials (AP) responsible for PP are generated only in the axons of the retinal ganglion cells that form the optic nerve. Pressure on the eyeball [6], as well as local mechanical deformations of the retinal layers [14, 15] lead to disturbances in the structure of Mueller glial cell membranes, which penetrate all layers of the retina, as well as photoreceptor membranes [16, 17, 18]. This changes the conductivity of voltage-dependent ion channels of membranes and the distribution of ion concentrations (Ca^**2+**^, Na^**+**^, Κ^**+**^ and Cl^**–**^) in the retinal tissues. Ionic currents in the conducting channels of membranes, in the cytoplasm and in the intercellular fluid correspond to polarization potentials, which, propagating along synaptic circuits in the layers of bipolar, horizontal and amacrine cells, reach the ganglion cells and activate them.

The background bioelectric activity of the retina is normally preserved in the dark and is manifested by the activity of ganglion cells (5-40 pulses/sec) [19] and the potential difference between the retina and the cornea (0.4-1.0 mV) [20]. Therefore, the retina is convenient for studying the mechanisms of interconnection between neurons of different layers and zones of the cerebral cortex. In addition, the phenomenon of phosphenes (PP and EP) makes it possible to obtain information on the participation of subcortical structures in the generation of AP in the VNS and on its connections with other sensory systems [21, 22].

When alternating current with a frequency of 4 Hz to 40 Hz is transmitted in the light between the Oz-Cz points (Fig. 1), the maximum EP rating is observed at frequencies of 16-17 Hz [23]. In the dark, the EP rating is halved and its maximum shifts to ∼12 Hz. Thus, the maxima of the EP ratings caused by alternating current in the dark and in the light are due to the resonant interaction of the current with the structures that generate the alpha and beta rhythms of the brain, respectively. The dark activity of the lateral geniculate body (LGB) is responsible for the alpha rhythm [24]. In addition to the retina and LGB, structures of the subcortex that regulate the movements of the eyeball (quadruple, hypothalamus, etc.) can also participate in the generation of the beta rhythm with open eyes [24].

**Fig. 1.**
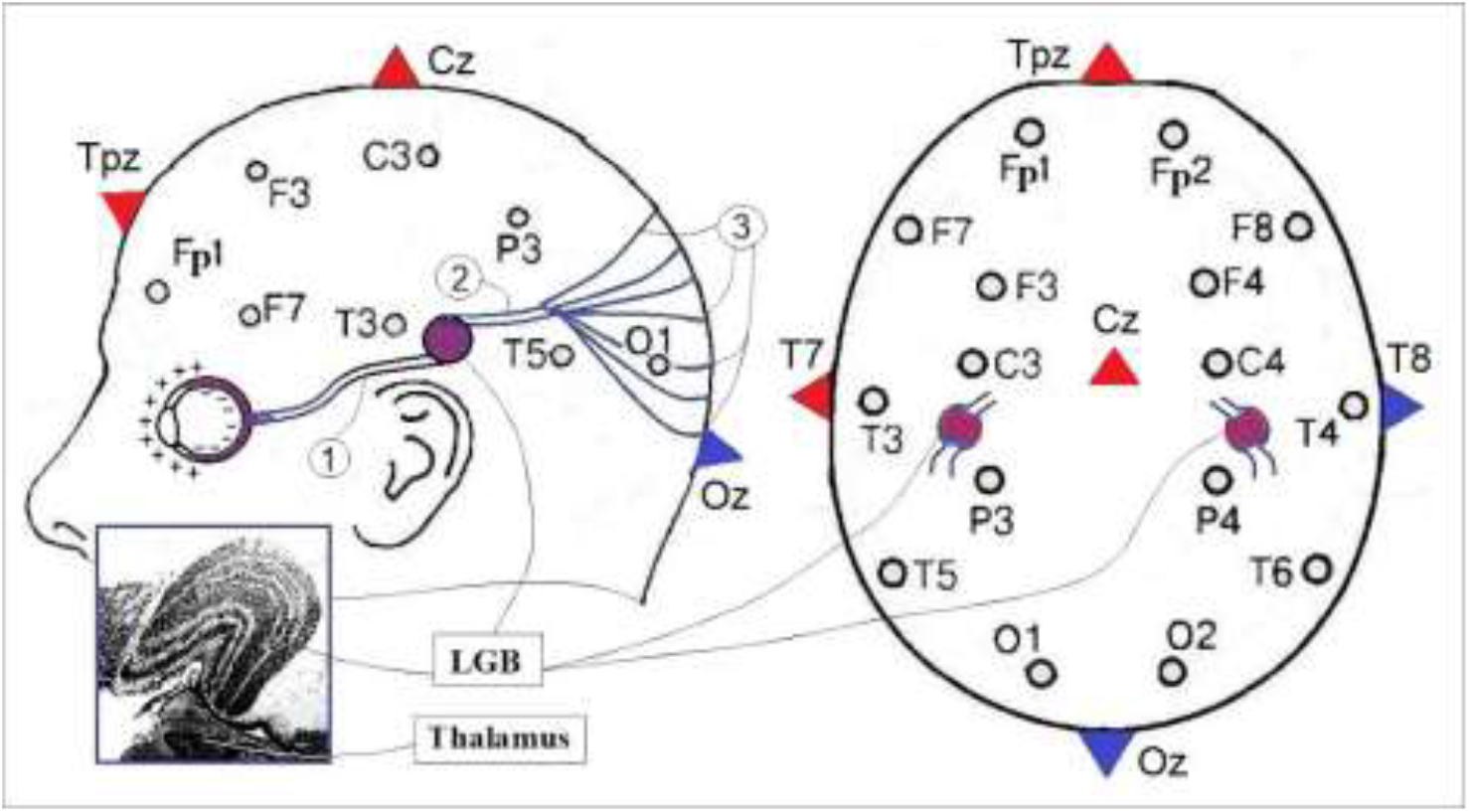
The standard scheme for the electrodes “10-20” on the projection of the brain and the layout of the structures of the visual apparatus in the brain: eye; 1 - optic nerve; LGB - lateral geniculate body, 2 - optic tract; 3 - fibers of the Graziole bundle. Inset LGB and thalamus. The triangles indicate the locations [23] of the anodes (red) and cathodes (blue).

From the analysis of the distributions of currents and fields when the EP is induced by an alternating current of 10 Hz [25] passing between the Fpz-Oz (F-O) and T7-T8 (T-T) electrodes (Fig. 1), it follows that the EP rating for the F-O option in ∼1.7 times higher than for T-T variant. The density of the induction current in the eyes for F-O is ∼2.6 times higher than for T-T. In this case, the values of the current density and field in the occipital lobe of the brain for the F-O variant are only ∼1.6 times higher than for the T-T variant. These ratios of ratings and currents between options F-O and T-T indicate that the EP induction by the current between points T7 and T8 is mainly due to the generation of AP in the right and left LGB.

It is known [6] that at high body temperature phosphenes appear more often and orgasm in men is sometimes accompanied by a sensation of a bright flash of white light. Ease of induction of phosphenes when the auditory system is excited [10, 26, 27, 28, 29] correlates with a high total proportion of visual synesthesia with four types of sound signals (Table 1). Cross-activation [30] of visual and auditory sensing indicates a physical relationship between LGB cells and cells of the medial geniculate body (MGB) [28, 29].

The order of the ganglion cells in the retina is reflected in the distribution of zones and cell layers in the LGB structure of the thalamus and is projected by the fibers of the Graziole bundle onto the visual cortex [6] (Fig. 1). Obviously, in this case, in the cellular structure of the thalamus and cortex, the subordination of cellular connections of the retina is reproduced in one form or another. This is evidenced by the high prevalence of synesthesia of grapheme-color modalities [30] (see Table 1), which may be due to the ease of cross-activation of adjacent LGB zones of the thalamus, as well as areas of the cortex VWFA (the grapheme processing) and V4 (a color processing) [30, 31, 32, 33, 34].

The onset of synesthesias is facilitated by the anatomical proximity of sensory systems and the retention of cross-neural connections in the early stages of perceptual brain development [31, 35]. These connections can also form as a consequence of the neuroplasticity of the cerebral cortex in response to its local damage or new perceptions [29, 30, 36]. In [31], a model of the mechanism of grapheme-color synesthesia is proposed, based on the principles of stochastic resonance and synchronization of increased neural noise in parallel modalities. Resonance can be established between oscillatory systems with close frequencies. Such systems can consist of coherent oscillatory LC circuits that model the current channels of axon membranes, including helical molecular structures [21, 37].

In [34], it is believed that synesthesias involving the color vision zone (V4) are associated with the network activity of several brain regions located in the sensory and motor regions, as well as in the higher-level zones of the parietal and frontal lobes. Many types of vision-induced synesthesias [3], as well as the strong influence of eye electrophysics on the electrical activity of the frontotemporal lobes of the brain, endow VNS with the function of the main conductor of the neurophysiology of cognitive processes [21]. This is also supported by phosphenes associated with synesthesias of vision with orgasm, hearing, tactile and thermal sensitivity [1, 2, 38]. Thus, studying the physics of pressure phosphenes will help to understand the mechanisms of cross-activation of neurons in different parts of the thalamus or areas of the cerebral cortex [39, 40] and the participation of subcortex structures in the neurophysiology of thinking and consciousness [21, 33, 34, 41, 42]. In practical terms, this information will be useful for optimizing the use of metameric therapy tools such as manual massage, acupuncture, exposure to low and high temperatures.

In this work, using EEG and ECG methods, we investigated the dependence of pressure phosphenes and bioelectric activity of the human brain and heart on the effect of light with different wavelengths and elevated temperatures on the eyes and brain.

## 2. Materials and methods

The studies involved 10 physically healthy men aged 32 to 72 years. In most of the experiments, the subjects were the authors of the article A.Kholmanskiy (AKN - N age), and A. Minakhin (AM60), a manual therapist with normal vision. AK – Doctor of Physical and Chemical Sciences, poet and painter [43, 44] had asymmetric myopia: -3D for the right (OD) eye and -3.75D for the left (OS). He was diagnosed with negative scotoma with visual field impairment OS by 14%, and OD by 7%.

A sauna with a temperature (T) of 60-90 °C was used for 10-15 minutes to heat the whole body in a humid and aromatized atmosphere, for which pure water or solutions of various essential oils were sprayed onto hot stones. Non-contact infrared thermometer measurements of body temperature in a sauna give 43 °C [45]. With manual massage, the skin temperature rises from ∼33 °C to ∼38 °C [46]. Steam condensation on the skin plays a significant role in the increase in body temperature in a sauna [47]. In the rest room, T of the body was 36.4 ± 0.2 °C. The hands were heated for 3-5 min in heated water (45-50 °C) or holding a polymer bottle (0.5 L) with heated water. The last version with a bottle (0.2 L) AK72 was used for local heating of both closed eyes for 1–2 min. PP was called after rest-relaxation (R) at room temperature (22-24 °C) for 10-15 minutes. The eyes were pressed for 10-30 seconds with the proximal phalanges of the thumbs of both hands or the thumb and forefinger of the right hand. The intensity of PP (I_P_) was assessed subjectively on a scale from 1 to 10 (1 – barely noticeable barely noticeable enlightenment of the field of view; 10 – bright white light in the entire field of view). The effect of white PPs on the LGB and visual cortex was simulated by irradiating the retina with the light of a wax candle, an alcohol lamp, and a white LED. In this case, a pupil-sized spot of light was focused on the eye with a glass lens with a diameter of 8 cm and a focal length of 20 cm.

Intraocular pressure (IOP) was measured using the Maklakov method and a Topcon CT-20 device with an accuracy of ±1 mmHg, as well as an HNT-7000 HUVITZ device (an accuracy of ±0.5 mmHg). The measurements showed that, when within the measurement error, the control IOP in OS and OD are equal and the pressure on healthy eyes AM60 leads to a decrease in IOP by the same amount, while in AK, the percentage of IOP reduction from pressure or eye heating in OS is usually higher than in OD (Table 2). These changes in IOP are consistent with the asymmetry of myopia and scotoma OS and OD.

**Table 2.**
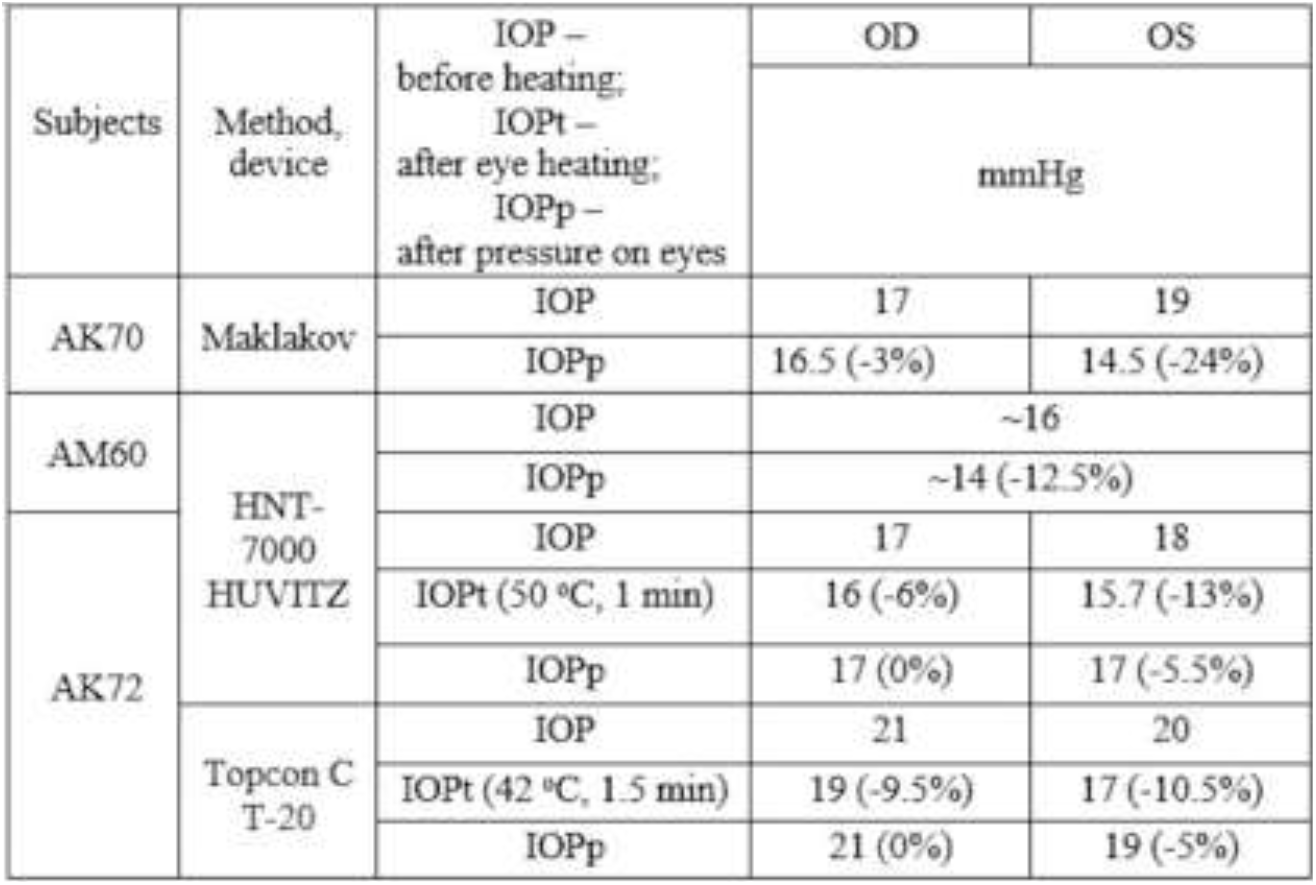
Intraocular pressure (IOP) versus external pressure and temperature. In parentheses, the percentage reduction from the reference value.

The bioelectrical activity of the heart and brain was recorded using an Encephalan-EEGR-19/26 (EG-1) electroencephalograph with a sampling rate of 250 Hz. The device allowed timing the oscillation frequencies (*ν*) from 0 to 100 Hz and the amplitudes of electric potentials (*V*) in the range from 0 to 0.5 mV. The data processing program of the device determined the average values of ν at intervals of 5 sec. From the standard 10–20 potential derivation, 4 points were selected on the left (F3, C3, P3, O1) and on the right (F4, C4, P4, O2), the reference electrodes were attached to the earlobes, the ground electrode to the forehead (Fig. 1).

For timing *ν* of the heart, electrodes were placed on the wrists at the point of a distinct pulse. The contact area of all electrodes was ∼0.7 cm^2^. With pressure on the eyes during registration of *ν*, a latex glove was put on the hand for electrical insulation. For the analysis, we used mainly frequency spectra EEG and ECG, which turned out to be more sensitive to the effects of pressure, heat and light on the eyes compared to the *V* spectra. For comparison, the distribution of changes in the *V* spectrum over the scalp from pressure on the eyes was also measured using a Compact-Neuro electroencephalograph (EG-2) with the imposition of 16 electrodes in a monopolar lead according to the standard “10-20” scheme (Fig. 1). The sampling rate was 500 Hz and the sweep speed was 30 mm/s.

## 3. Results

In all 10 subjects, after the sauna, bright white PPs were easily evoked by pressure on both eyes. In most experiments, the glow began to flare up from 3 seconds, after 7-10 seconds it reached a maximum with I_p_ ∼ 7-10 and faded away by 15-20 seconds. At the end of the pressure on the eyes, blue-violet dots could appear against the background of weak white light. AK49 sometimes felt the same phosphenes in the morning after waking up with light pressure on both eyes. Manual massage of the cervical and thoracic spine, performed by AM60 to himself and three subjects, led to a relief in the induction of PP with I_p_ ∼ 8. The PP intensity was markedly reduced with alternate pressure on only one eye.. In AK70 and AK72, PP induction was facilitated by heating the hands in water with T ∼ 45-50 °C. On the other hand, after heating under the same conditions, the feet to the ankle with a temperature of 50-60 °C did not develop PP. There was no positive effect from heating the hands and eyes with dry plastic bottles containing water with T ∼ 45-50 °C. When the sauna air was saturated with water vapor or calendula ester vapors, I_p_ of PP did not change, but mustard mint vapors significantly reduced I_p_.

### 3.1. Informative value of EEG and ECG spectra

Frequency (*ν*) and amplitude (*V*) spectra of EEG and ECG are given in in Appendix (Figs. 2-7). The principle of coding information by receptors is based on the variations in the repetition rate of AP ([1, 6, 20]. ν-Spectra are also sensitive to changes in the local cerebral circulation of neurons [48], which is modulated by electrical oscillations of blood electrolytes generated by cardiomyocytes [39]. Therefore, the frequency spectra of the ECG and EEG (Figs. 2) in our case best reflect the bioelectric reactions of the brain to external factors. The informative value of *ν* spectra of ECG and EEG, obtained by the EG-1 method with pressure on the eyes with a hand without a glove, sharply decreases in all areas of the brain and especially at points F8, F7 (Figs. 2 and 6). This may be due to the shunting of currents in the brain when the electrode on the wrist approaches the ground electrode on the forehead.

**Fig. 2.**
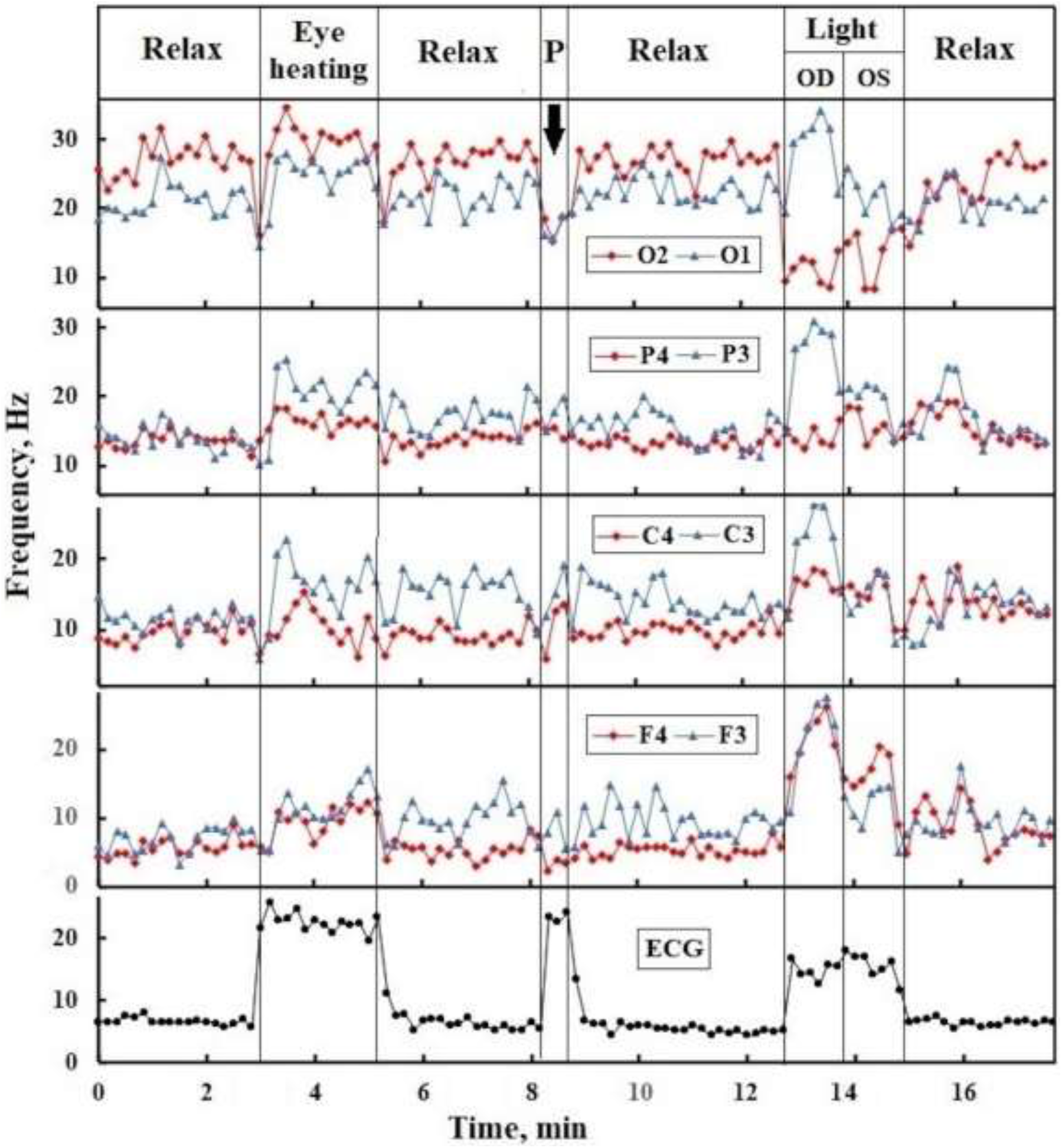
Dependence of the frequency *ν*-spectra of EEG and ECG AK72 on heating of both eyes by objects with a temperature of ∼50 °C, pressure on the eyes with gloved fingers (**P**) alternating wax candle lighting of the right (OD) and left (OS) eyes AK72.

**Fig.3.**
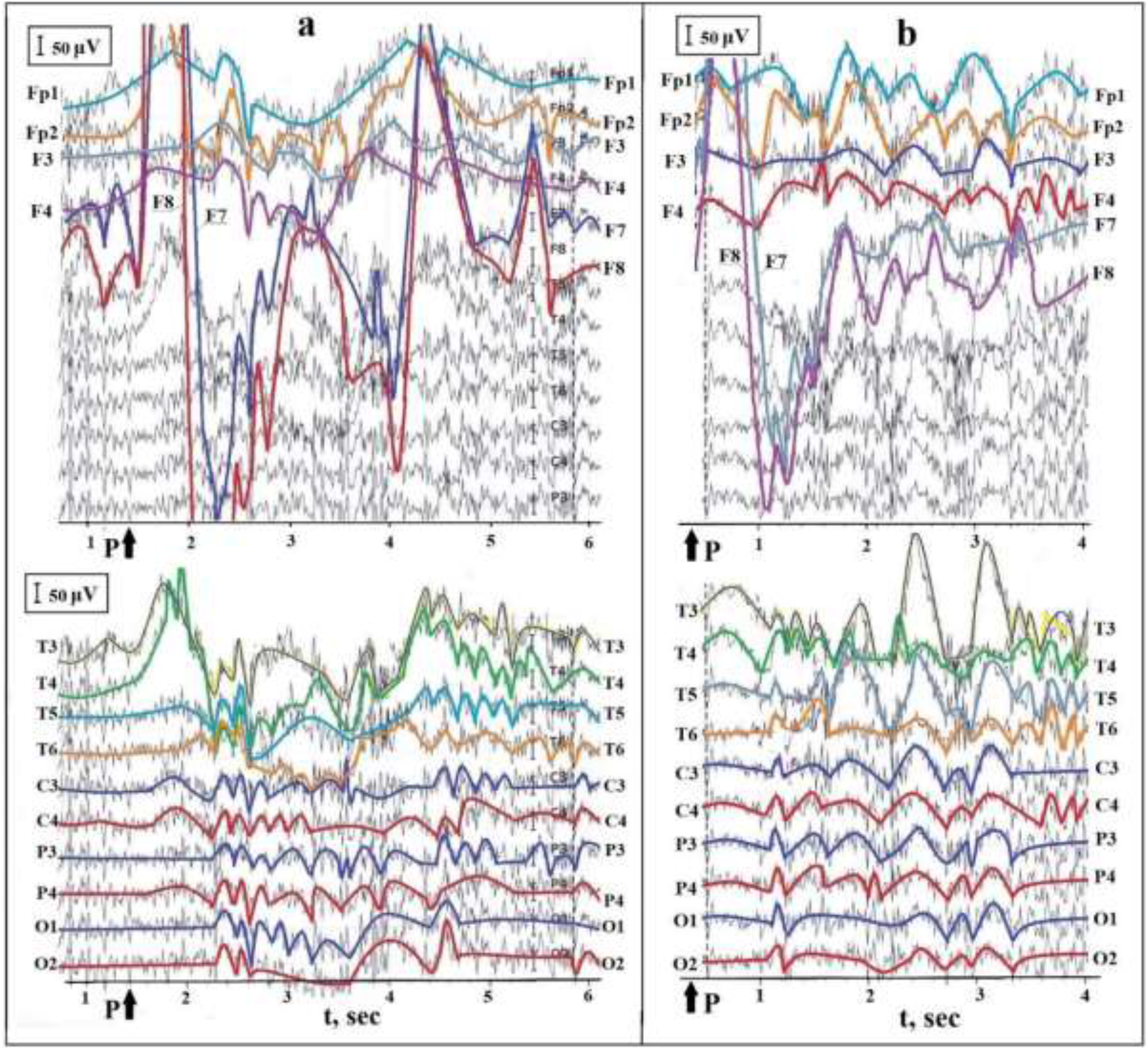
Dependence of *V*-spectra of AK70 on the pressure (**P**) on the eyes with fingers without gloves (**a**) and a repeat of the experiment after ∼2 min (**b**). Colored lines bend around the low-frequency *V* oscillations at each contact point.

**Fig. 4.**
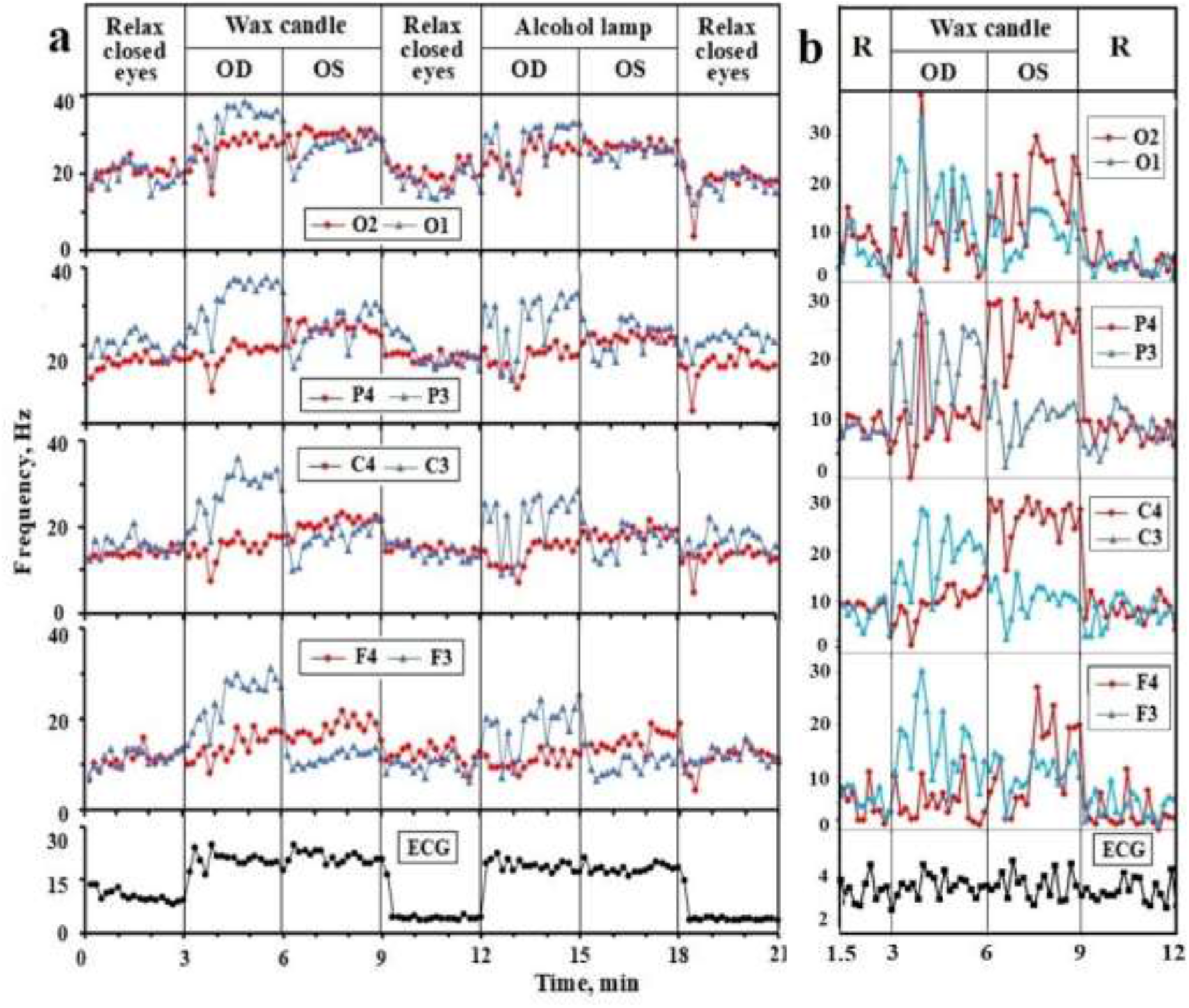
Dependence of *ν*-spectra of EEG and ECG (**a**) AK70 and (**b**) AM on irradiation of the eyes with the light of a wax candle and an alcohol lamp.

**Fig. 5.**
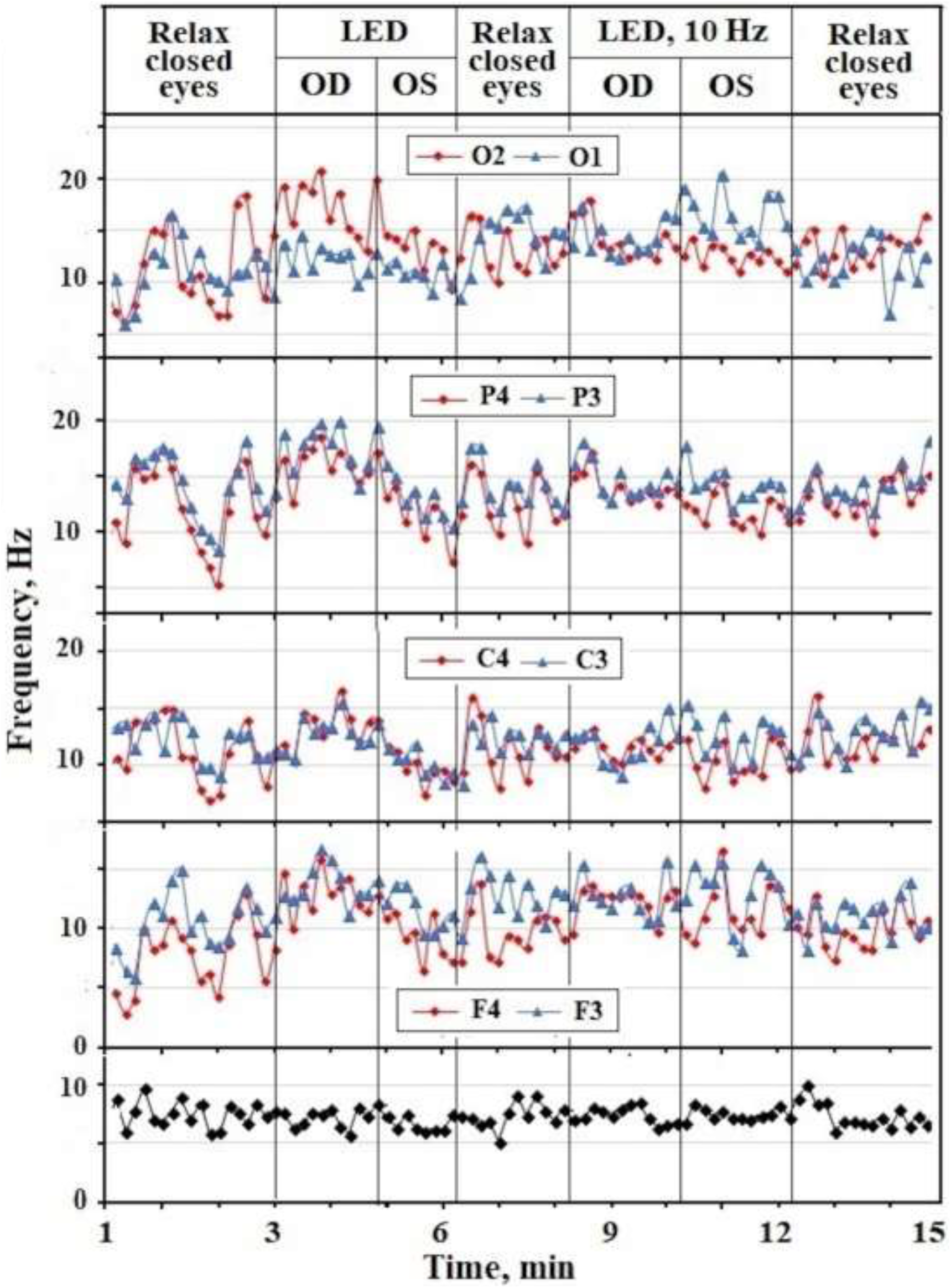
Dependence of *ν*-spectra of EEG and ECG AK70 on irradiation with continuous and flashing (10 Hz) LED light.

**Fig. 6.**
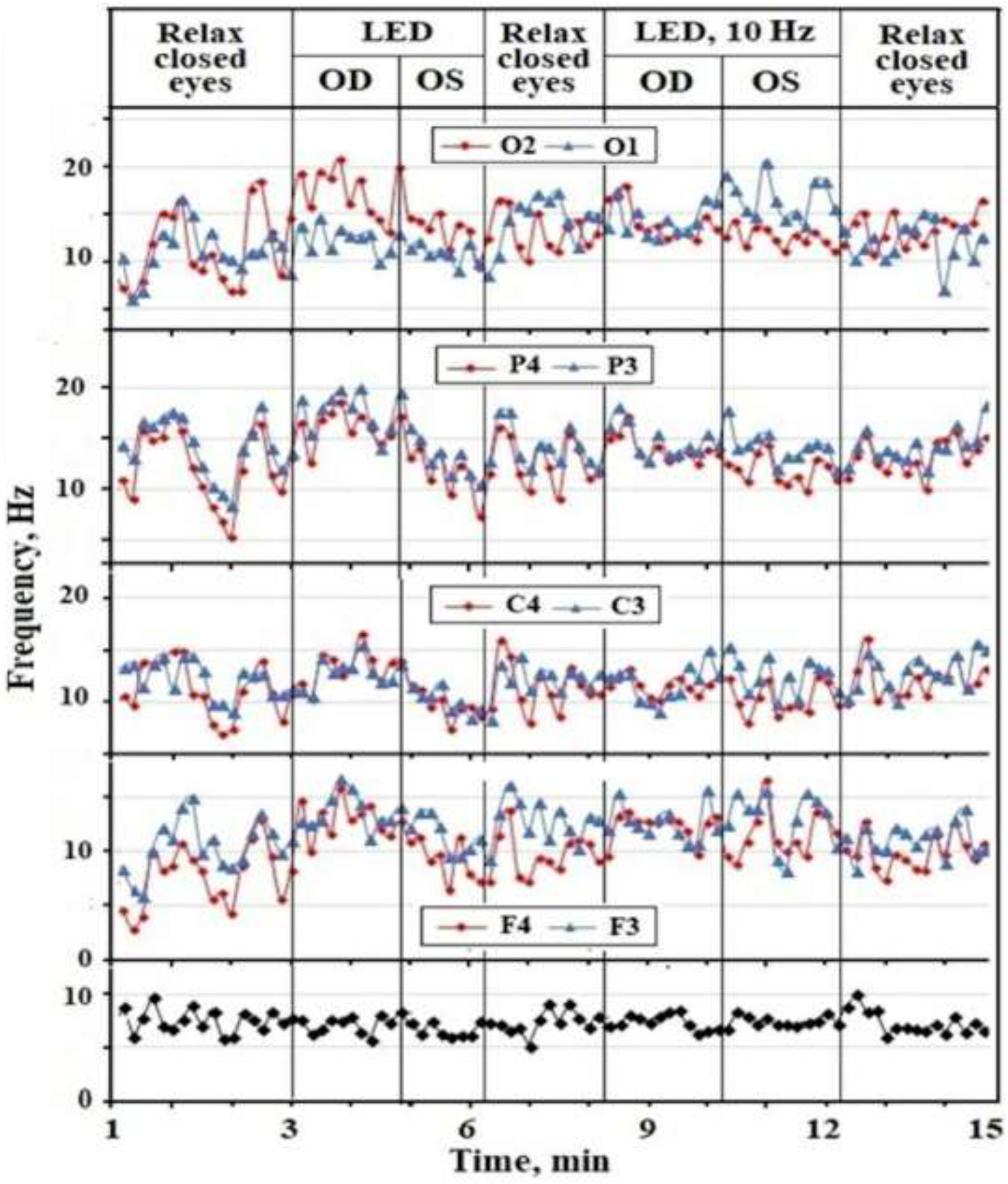
Dependence of *ν*-spectra of EEG AK70 on pressure (**P**) on both eyes (OU) and alternately on the right (OD) and left (OS) eyes with fingers in a latex glove (**a**) and without it (**b**).

**Fig. 7.**
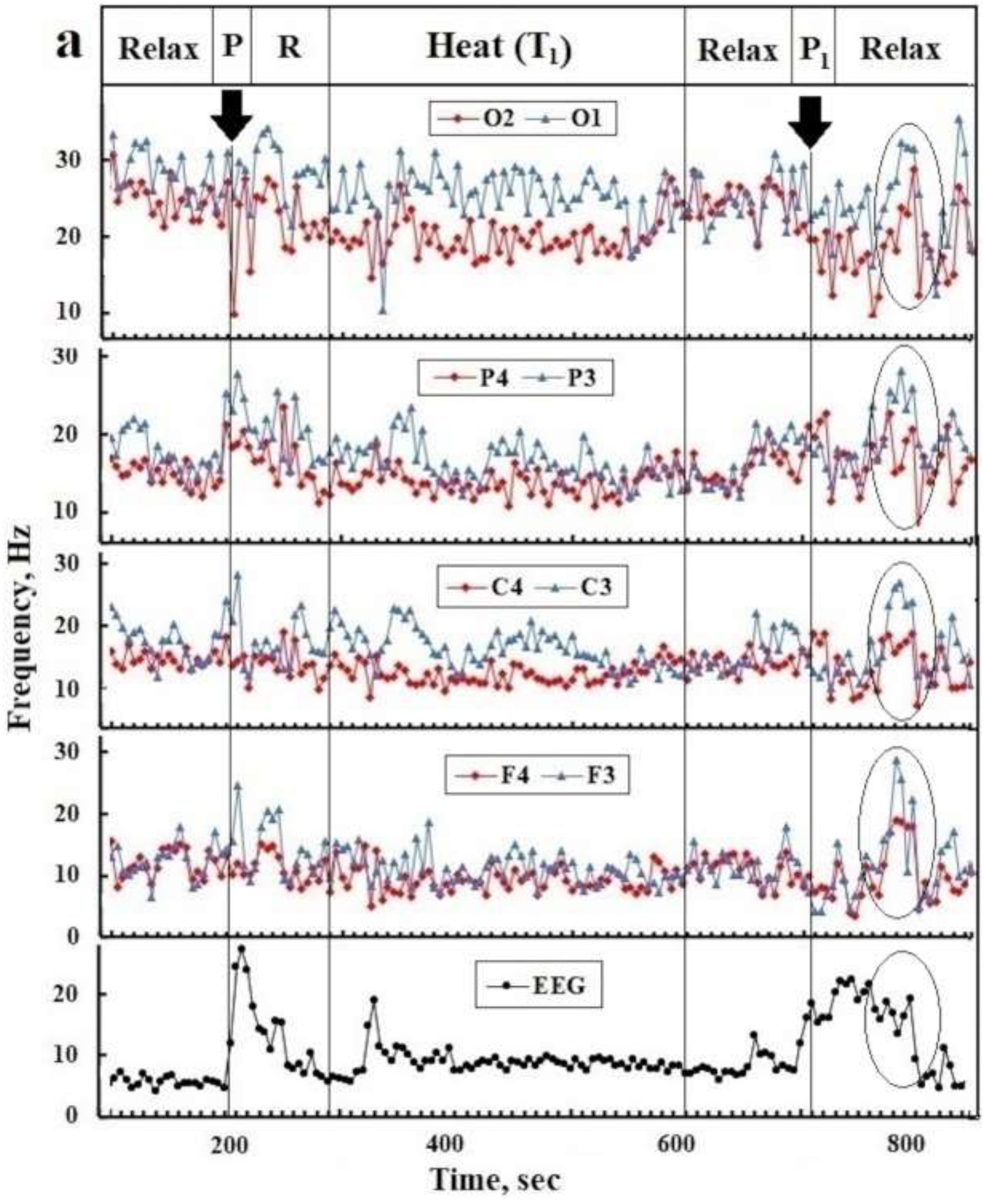
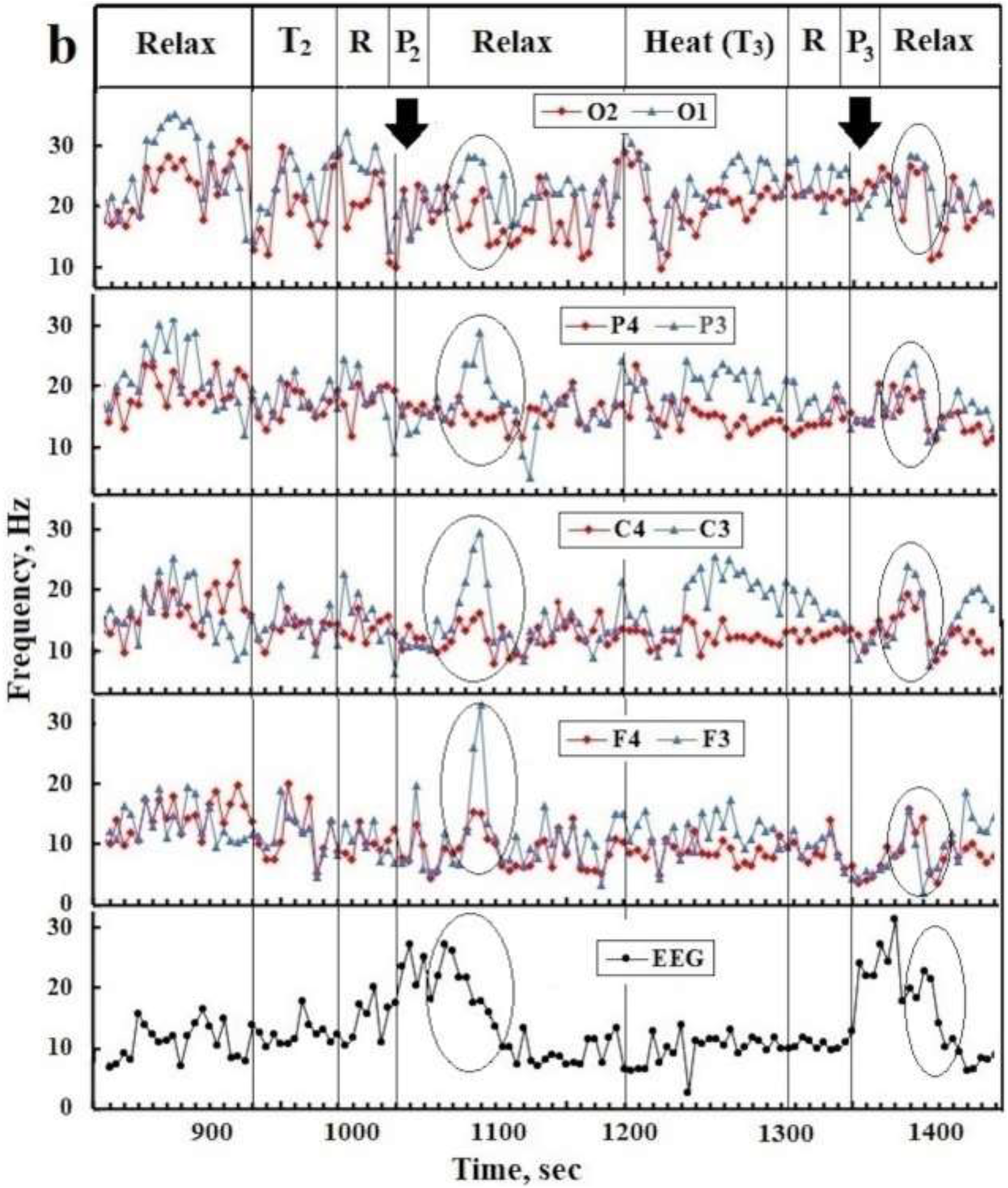
Dependence of *ν*-spectra of EEG AK72 on pressure on both eyes for 20 seconds with gloved fingers before (**P**) and after warming up the hands (**P**_**1**_, **P**_**2**_, **P**_**3**_); (**a**) dry objects with T_1_ ∼ 50 ° C; (**b**) in water with T_2_ ∼ 48 °C and T_3_ ∼ 43 °C. R - relaxation with closed eyes. Ovals mark the frequency deviations caused by pressure.

### 3.2. Asymmetry and kinetics of reactions of the visual system to light and pressure

It is known [49, 50, 51, 52] that the area of the nasal half of the eyes and the number of photorecentors in it are greater than in the temporal half. Since the axons of neurons in the nasal part of the retina pass in the chiasm into the optic tract of the opposite hemisphere, then normally, when the eyes are irradiated with light in all ν EEG spectra, the signals from the contralateral hemisphere are higher than from the lateral hemisphere (Fig. 4b). In AK70, this effect is enhanced for OD and almost absent in OS (Figs. 2 and 4a). This asymmetry of VNS in AK70 can be explained by the specificity of creative mental work [43, 44, 53] and the difference in the physiology of OS and OD in AK70, which manifests itself in scotoma, myopia, and IOP (Table 2). The contralateral effect depends on the spectrum of the light source: it is most pronounced for a wax candle, decreases for a spirit lamp flame (Figs. 2 and 4) and is practically absent for an LED (Fig. 5). The first two thermal light sources have emission maxima at ∼1000 nm, and LED in the 400-600 nm region. Obviously, the sensitivity of the nasal and temporal halves of the retina for LED light is the same, while for IR light it predominates in the nasal half.

From the *V* spectra obtained with the EG-2 (Fig. 3) it follows that the pressure on both eyes causes an oscillation in the points F7 and F8 closest to the eyes with a period of ∼2 sec and an initial value of *V* ∼ 1.5 mV. This oscillation decays after two periods, and with repeated pressure on the eyes, its amplitude decreases to ∼0.3 mV and decays twice as fast. Note that in the *V* spectra the difference in oscillations in pairs of contralateral points is weakly expressed.

The *V* values are comparable to the magnitude of the potential on the cornea of the eye and the onset of oscillations at points F7 and F8 is synchronized with the application of pressure. Given this and the orientation of the eye dipole line to points F7 and F8, the short-lived oscillation *V* can be attributed to electrooculogram signals caused by deformation of the eye muscles and the eyeball dipole (Fig. 1) [20]. The directions to points Fp1 and Fp2 are orthogonal to the eye dipoles, and therefore the oscillations of *V* at these points have an amplitude of ∼0.2 mV. On the other hand, the amplitudes *V* in the pairs of points (T3, T4) and (T5, T6) reach values of 0.3-0.4 mV with a delay of ∼ 0.7 s from the beginning of pressure on the eyes. Hence, it follows that the likely stimulus for oscillations at these points is the redistribution of charges in the LGB layers, initiated by currents from retinal deformation. With the same delay, but with a smaller amplitude, *V* oscillations are excited in pairs of points (C3, C4), and in pairs (P3, P4), (O1, O2), the delay increases to ∼1 sec (Fig. 3a) and is consistent with the delay PP start flaring up. The decrease in the frequency-amplitude characteristics of EEG signals with repeated pressure on the eyes can be explained by a decrease in the concentration of charges in the layered tissues of the retina and LGB, due to their recombination caused by the first pressure. In [21], such structures are modeled by capacitor systems as part of neural networks.

### 3.3. Influence of heating of hands and eyes on the frequency spectra of EEG and ECG

The frequency of the ECG spectrum increases ∼4 times with heating and pressure on the eyes, while synchronous increases in the frequencies of the EEG spectra are observed only during heating and differ for different points (Fig. 2). Pressure on the eyes after heating them, as well as without heating, did not cause the sensation of PP (Ip ∼ 0), however, it led to a change in the background spectra of EEG and ECG frequencies (Figs. 2 and 6). The same result was given by a control experiment of pressure on the eyes before warming the hands (Fig. 7a). The pressure on the eyes after heating the hands with plastic bottles with water with T_1_ ∼ 50 °C caused a weak glow over the entire field with an increase to the equator line (Ip ∼ 1-2). The glow flared up for 5-7 seconds and quickly died out after the pressure on the eyes was stopped, and ∼1 min after that, a frequency jump was observed in all EEG spectra at all points and especially in the left hemisphere. The disturbance of the ν-spectrum of the ECG was synchronized with it. These disturbances of the ECG and EEG spectra are highlighted by ovals in Fig. 7a. After heating both hands in water for 1 min (T_2_ ∼ 48 °C), the intensity of PP increased to 5-6; when the hands were reheated after 2 min for 2 min in water (T_3_ ∼ 43 °C), pressure on the eyes did not cause PP (Fig. 7b). Note that changes in the ν-spectra of ECG and EEG caused by pressure on the eyes after heating the hands in water with T_2_ and T_3_ were more noticeable than after heating with dry objects with T_1_ (Fig. 7b).

## 4. Discussion

The time of AP propagation from the retina along the VNS correlates with the delay time of the PP, and the bioelectric activity of the retina and LGB is consistently manifested in the EEG spectra by potentials at points (F7, F8), (T3, T4), (C3, C4) and (O2, O1). The dynamics of the evoked potentials in each pair of points can be modeled by a rapidly decaying sinusoidal signal generated by the charge recombination current, first between the retinal layers, then the LGB layers [21, 39]. In the layers of the retina, these processes are initiated by the mechanical deformation of its tissues, and the bioelectrical activity in the LGB layers is caused by the AP flows emanating from the retina and corresponding to the PP. Since direct eye heating does not stimulate PP generation, this effect should be realized at the level of LGB neurophysiology. The presence of neural connections between LGB and MGB (synesthesia of vision and hear) indicates the possibility of a neurophysiological relationship between LGB and the nuclei of the hypothalamus and thalamus, which are responsible for thermoreception [1, 54, 55]. The latter include the thalamic nuclei closest to the LGB: the ventral medial nucleus (VM), the ventral medial nucleus (VM), the ventral posterior medial nucleus (VPM), and the pulvinar nuclei (PUL).

In the course of evolution, thermoreceptors developed as modifications of mechanoreceptors, primarily in the skin of the fingers, palms of the hands, and in the cornea of the eyes [1]. The molecular mechanisms of pain and thermoreceptors are not yet fully understood [54, 55]. It is known ([1, 2, 6, 56, 57] that cold and heat receptors at T≥42 °C begin to work as pain receptors. It is believed that at these temperatures, the cold receptor mediator menthol [58; 59] will also affect the conductance of other protein ion channels of receptors of the TRPA, TRPV, and TRPM families [55, 56, 57]). Menthol, forming a hydrogen bond with proteins, causes the opening of ionic Ca^2+^ channels and the flow of Ca^2+^ across the membrane of receptor cells leads to their inactivation and loss of the ability to respond to stimuli [55, 58, 59]. The action of menthol, which contains ∼40% in mustard mint, can explain the decrease in the thermal effect of PP stimulation when mint is added to the sauna at temperatures 60-90 °C. The lack of a thermal effect upon direct heating of the eyes and cornea is apparently due to the lack of convergence with the VNS centers of the trigeminal nerve fibers, which innervate the corneal heat and pain receptors.

Due to its incompressibility, water in the vitreous humor and intercellular fluid of the retina evenly distributes the action of the pressure force over all retinal tissues tightly adjacent to the elastic and inextensible sclera of the eye. The absence of PP at the sites of retinal detachment from the choroid feeding it [60] indicates the key role of the physiological fluids of the retina and mechanical resistance from the sclera in the process of PP generation. There are no special moisture receptors in the epidermis of the palms and fingers. Their absence is compensated for by the responses of tactile mechanoreceptors to water pressure and thermoreceptors to the thermal effects of water condensation and evaporation [47, 61]. Hydrophilicity of the skin implies the formation of hydrogen bonds of water with epidermal molecules when hands are immersed in water. As a result of these bonds, the efficiency of resonant heat transfer from water to the blood and epidermal tissues, and therefore to the thermoreceptors, increases. Heat transfer when the hands are heated with a plastic bottle with heated water will be significantly worse, and this explains the absence of the thermal effect of PP stimulation in this case.

In addition to water entering the hydration shells of ions and associated with biomolecules, a certain proportion of free water is also present in the blood plasma and in the intercellular fluid [62, 63]. Anomalies of its physical properties in the range from ∼25 °C to ∼ 42 °C are determined by the dynamics of hydrogen bonds in clusters of various compositions [64, 65, 66]. These features of the thermodynamics of water and its hydrogen bonds with the proteins of the conductive channels of cell membranes can underlie the differentiation of the mechanisms of operation of cold, heat and pain receptors in the epidermis. Apparently, the leveling of these features of water thermodynamics at T> 42 °C leads to the observed unification of the mechanism of the pain and thermoreceptors.

## 5. Conclusion

The generation of pressure phosphenes is based on the mechanical activation of bioelectric processes in the layers of the retina, which normally occur in it in dark conditions. At the same time, the efficiency of the generation of action potentials in ganglion cells depends on the state of the physiology of the eyes and its adaptation to the professional specificity of the use of vision. Stimulation of the process of generation of pressure phosphenes by preliminary heating of the hands in water or manual massage of the cervical and thoracic spine indicates the presence of convergence between LGB neurons and neurons of the thalamic nuclei, which control the somatosensory of the body.

## Ethics approval and consent to participate

Not applicable.

## Consent for publication

Not applicable.

## Availability of data and material

Data sharing not applicable to this narrative review

## Competing interests

We have no competing interest

## Funding statement

None

## Author Contributions

Conceptualization and methodology: AK, AM; Software, formal analysis, investigation: AΚ, EK; writing, AK.

## Notes

https://www.longdom.org/open-access/thermal-stimulation-neurophysiology-of-pressurephosphenes-80515.html

### Competing Interest Statement

The authors have declared no competing interest.

### Summary of Updates

Removed 2 pictures and edited the text by adding links

## References

1. Smith, C.U.M., (2000) Biology of Sensory Systems. Chichester: John Wiley. New York.

2. Manivannan, M., Suresh, P. K. (2012). On the somatosensation of vision. Ann. Neurosci. 19, 31–39. 10.5214/ans.0972.7531.180409,

3. Day, S.A. (2020) Demographic aspects of synesthesia. http://www.daysyn.com/Types-of-Syn.html

4. Kholmanskiy, A., Zaytseva, N. (2018) Chiral Factor of Circadian Rhythm of Human physiology, Int. J. Res. Pharm. Biosci. 5(4) 6–10. https://www.ijrpb.org/papers/v5-i4/2.pdf.

5. Wirz-Justice, A., Skene, D.J., Münch, M. (2020) The relevance of daylight for humans, Biochemical Pharmacology, 114304. https://doi.org/10.1016/j.bcp.2020.114304.

6. Human Physiology (1989) Ed. R.F. Schmidt and G. Thews. Springer-Verlag, Berlin-Heidelberg-New York.

7. Medanic, M., Gillette, M.U. (1993) Suprachiasmatic circadian pacemaker of the rat shows two windows of sensitivity to neuropeptide Y in vitro. Brain Res. 620 (2), 281–6.

8. Grüsser, O.-J., Hagner, M. (1990) On the history of deformation phosphenes and the idea of internal light generated in the eye for the purpose of vision. Documenta Ophthalmol. 74: 57–85. 10.1007/BF00165665.

9. Lövsund, P., Öberg, P. Å., Nilsson, S. E. G. (1980) Magneto- and electrophosphenes: a comparative study, Medical & Biological Engineering & Computing, 18(6) 758–764. https://link.springer.com/article/10.1007/BF024419023.

10. Nair, A., Brang, D. (2019) Inducing synesthesia in non-synesthetes: Short-term visual deprivation facilitates auditory-evoked visual percepts. Conscious Cogn. 70, 70–79. 10.1016/j.concog.2019.02.006

11. Sappington, R. M., Sidorova, T., Ward, N. J., Chakravarthy, R., Ho, K. W., Calkins, D. J. (2015) Activation of transient receptor potential vanilloid-1 (TRPV1) influences how retinal ganglion cell neurons respond to pressure-related stress. Channels, 9, 102–113. 10.1080/19336950.2015.1009272

12. Höfling, L., Oesterle, J., Berens, P. Zeck (2020) Probing and predicting ganglion cell responses to smooth electrical stimulation in healthy and blind mouse retina, Scientific Reports 10(1) 10.1038/s41598-020-61899-y

13. Grüsser, O.-J., Grüsser-Cornehls, U., Kusel, R., Przybyszewski, A.W. (1989) Responses of retinal ganglion cells to eyeball deformation: A neurophysiological basis for “pressure phosphenes” Vision Res. 29, 181–194, https://doi.org/10.1016/0042-6989(89)90123-5

14. Marrese, M., Lonardoni, D., Boi, F., Van Hoorn, H., Maccione, A., Zordan, S., Iannuzzi, D., Berdondini, L. (2019) Investigating the Effects of Mechanical Stimulation on Retinal Ganglion Cell Spontaneous Spiking Activity, Front. Neurosci., https://doi.org/10.3389/fnins.2019.01023

15. Rountree, C. M., Meng, C., Troy, J. B., Saggere, L. (2018). Mechanical stimulation of the retina: therapeutic feasibility and cellular mechanism. IEEE Trans. Neural Syst. Rehabil. Eng. 26, 1075–1083. 10.1109/TNSRE.2018.2822322

16. de Melo Reis, R.A., Marques Ventura, A.L., Schitine, C.S., Fialho de Mello, M.C., de Mello F.G. (2008) Müller glia as an active compartment modulating nervous activity in the vertebrate retina: neurotransmitters and trophic factors. Neurochem. Res. 33:1466–1474. 10.1007/s11064-008-9604-1

17. Lindqvist, N., Liu, Q., Zajadacz, J., Franze, K., Reichenbach, A. (2010). Retinal glial (Müller) cells: sensing and responding to tissue stretch. Investig. Opthalmol. Vis. Sci. 51, 1683–1690. https://doi.org/10.1167/iovs.09-4159

18. Agte, S., Pannicke, T., Ulbricht, E., Reichenbach, A., Bringmann, A. (2017). Two different mechanosensitive calcium responses in Müller glial cells of the guinea pig retina: differential dependence on purinergic receptor signaling. Glia, 65, 62–74. https://doi.org/10.1002/glia.23054

19. Hubel D.H. (1995) Eye, Brain, and Vision, Scientific American Library Series. New York.

20. Malmivuo, J., Plonsey, R., (1995) Bioelectromagnetism - Principles and Applications of Bioelectric and Biomagnetic Fields, New York, 576–587, 10.1093/acprof:oso/9780195058239.001.0001

21. Kholmanskiy, A. (2006) Modeling of brain physics. Mathematical morphology. Electronic mathematical and Medico-biological journal. 5 (4). https://www.preprints.org/manuscript/201906.0188/v1

22. Baysal, V., Yilmaz, E. (2020) Effects of electromagnetic induction on vibrational resonance in single neurons and neuronal networks, Physica A: Statistical Mechanics and its Applications, 537, article id. 122733. 10.1016/j.physa.2019.12273312.

23. Kanai, R., Chaieb, L., Antal, A., Walsh, V., Paulus, W. (2008) Frequency-Dependent Electrical Stimulation of the Visual Cortex, Current Biology, 18, 1839–1843, 10.1016/j.cub.2008.10.027

24. Jensen O., Spaak E., Zumer J.M. (2019) Human Brain Oscillations: From Physiological Mechanisms to Analysis and Cognition. In: Supek S., Aine C. (eds) Magnetoencephalography. Springer, Cham. 471–517. https://doi.org/10.1007/978-3-030-00087-5_17

25. Indahlastari, A., Kasinadhuni, A.K., Saar, C., Castellano, K., Mousa, B., Chauhan, M., Mareci, T.H., Sadleir, R.J. (2018) Methods to Compare Predicted and Observed Phosphene Experience in tACS Subjects, Neural lasticity, 8525706, 10 pages, https://doi.org/10.1155/2018/8525706

26. Page, N.G.R., Bolger, J.P., Sanders, M.D. (1982) Auditory evoked phosphenes in optic nerve disease, J. Neurology, Neurosurgery and Psychiatry. 45(1): 7–12. 10.1136/jnnp.45.1.7

27. Lessell S., Cohen M.M. (1979) Phosphenes induced by sound. Neurology. 29(11) 1524–1526. 10.1212/wnl.29.11.1524

28. Brang, D., Towle, V.L., Suzuki, S., Hillyard, S.A., Di Tusa, S., Dai, Z., Tao, J., Wu, S., Grabowecky, M. (2015) Peripheral sounds rapidly activate visual cortex: evidence from electrocorticography, J. Neurophysiol. 114(5) 3023–3028. 10.1152/jn.00728.2015

29. Roe, A.W., Garraghty, P.E., Esguerra, M., Sur, M. (1993) Experimentally induced visual projections to the auditory thalamus in ferrets: Evidence for a W cell pathway, J. Comp. Neurol. 334(2): 263. https://doi.org/10.1002/cne.903340208

30. Hubbard, E.M., Brang, D., Ramachandran V.S. (2011). The cross-activation theory at 10. J. Neuropsychol. 5(2), 152–177. 10.1111/j.1748-6653.2011.02014.x

31. Lalwani P., Brang D. (2019) Stochastic resonance model of synesthesia. Philos. Trans. R Soc. Lond. B Biol. Sci. 374(1787):20190029. 10.1098/rstb.2019.0029

32. Rouw, R, Scholte, H.S. (2007) Increased structural connectivity in grapheme–color synesthesia. Nat. Neurosci. 10, 792–797. doi:10.1038/nn1906

33. Ramachandran, V.S., Hubbard, E.M. (2001) Synaesthesia – a window into perception, thought and language. J. Conscious. Stud. 8, 3–34. ISI, Google Scholar

34. Rouw, R, Scholte, H.S., Colizoli O.J. (2011) Brain areas involved in synaesthesia: a review. J. Neuropsychol. 5(2) 214–42. 10.1111/j.1748-6653.2011.02006.x

35. Maurer, D, Ghloum, J.K., Gibson, L.C., Watson, M.R., Chen, L.M., Akins, K., Enns, J.T., Hensch, T.K., Werker, J.F. (2020) Reduced perceptual narrowing in synesthesia. Proc. Natl. Acad. Sci. U S A. 117(18):10089–10096. 10.1073/pnas.1914668117

36. Sur, M., Garraghty, P. E., Roe, A. W. (1989) Experimentally-induced visual projections into auditory thalamus and cortex, Science, 242, 1437—1441.

37. Kholmanskiy, A. (2019) Dialectic of Homochirality. Preprints, 2019060012. https://www.preprints.org/manuscript/201906.0012/v1

38. Gaglianese, A., Branco, M.P., Groen, I.I.A. Benson, N.C., Vansteensel, M.J., Murray, M.M., Petridou, N., Ramsey, N.F. (2020) Electrocorticography Evidence of Tactile Responses in Visual Cortices, Brain Topogr. 33(5) 559–570, 10.1007/s10548-020-00783-4

39. Kholmanskiy, A.S., Minakhin, A.A. (2018) Interconnection of electrical oscillations of the heart and brain. Bull. St.-P. State Univ. Med. 13(2) 117–135. https://dspace.spbu.ru/bitstream/11701/10429/1/01-Kholmansky.pdf

40. Snyder, A. C., Issar, D., Smith, M.A. (2018) What does scalp electroencephalogram coherence tell us about long-range cortical networks? European J. Neuroscience, 48(7) 2466–2481. https://doi.org/10.1111/ejn.13840

41. Hubbard, E.M., Ramachandran, V.S. (2005). Neurocognitive mechanisms of synesthesia. Neuron, 48 (3): 509–520. 10.1016/j.neuron.2005.10.012. PMID 16269367

42. Vandewalle, G., Maquet, P., Dijk, D.-Jan. (2009) Light as a modulator of cognitive brain function, Trends Cogn. Sci. 13(10):429–38. 10.1016/j.tics.2009.07.004

43. Kholmanskiy, A. (1998) The Way of Salvation. 133, https://search.rsl.ru/ru/record/01000632964

44. Kholmanskiy, A. (2010) Flowers of Light, https://search.rsl.ru/ru/record/01004733425

45. Hermans, L.F.J (Jo), Vesala, T., (2007) The sauna – revisited, Europhysics News 38(6) 32–32. https://doi.org/10.1051/epn:200702

46. Kholmanskiy, A.S., Minakhin, A.A. (2011) Physical factors of asymmetry of biomechanics of the musculoskeletal system, Mathematical morphology. Electronic mathematical and Medico-biological journal. 10(3), http://sgma.alpha-design.ru/MMORPH/N-32-html/holmanskiy/holmanskiy.htm

47. Zech, M., Bösel, S., Tuthorn, M., Benesch, M., Dubbert, M., Cuntz, M., Glaser, B. (2015): Sauna, sweat and science – quantifying the proportion of condensation water versus sweat using a stable water isotope (2H/1Hand 18O/16O) tracer experiment, Isotopes in Environmental and Health Studies, 10.1080/10256016.2015.1057136, http://dx.doi.org/10.1080/10256016.2015.1057136

48. Fokin V.F., Ponomareva N.V. Energetic physiology of the brain, Moscow, Antidor, in Russian, 2003. 288.

49. Bogoslovsky A.I. (1962) General pathology of the visual field. In the book Multivolume Guide to Eye Diseases, V. 1, Moscow., 492-502, (In Russian).

50. Curcio, C.A., Allen, K.A. (1990) Topography of ganglion cells in human retina, J. Comp. Neurol. 300. 5–25. 10.1002/cne.903000103

51. 52. Sjöstrand, J., Olsson, V., Popovic, Z., Conradi, N. (1999) Quantitative estimations of foveal and extra-foveal retinal circuitry in humans, Vision Res. 39(18), 2987–2998, https://doi.org/10.1016/S0042-6989(99)00030-9

52. Muthuswamy A., Chen H., Ying Hu Y. et al. (2020) Mammalian Retinal Cell Quantificationr Toxicologic Pathology 49(210): 019262332097637, 10.1177/0192623320976375

53. Dashchinskaya, T.N., Dashchinskiy V.E., Kholmanskiy A.S. (2012) Sociocultural and psychophysical foundations of artistic creativity, Modern problems of science and education..2, http://science-education.ru/ru/article/view?id=5918.

54. Filingeri, D. (2016) Neurophysiology of Skin Thermal Sensations, Compr. Physiol. 6(3). 1279–1294. https://doi.org/10.1002/cphy.c150040

55. Medvedev, A.A., Sokolova, L.V. (2019) Features and Mechanisms of Temperature Sensitivity (Review). J. Med. Bio. Res. 7(1) 92–105. 10.17238/issn2542-1298.2019.7.1.92

56. Basbaum, A.I., Bautista, D.M., Scherrer, G., Julius, D. (2009) Cellular and Molecular Mechanisms of Pain, Cell. 139(2): 267–284. 10.1016/j.cell.2009.09.028

57. Fernández-Carvajal, A., Fernández-Ballester, G., Devesa, I. González-Ros, J. M. Ferrer-Montiel, (2012) New Strategies to Develop Novel Pain Therapies: Addressing Thermoreceptors from Different Points of View, Pharmaceuticals, 5, 16–48; 10.3390/ph5010016

58. Kozyreva, T.V., Tkachenko, E.Ya. (2008) Effect of menthol on human temperature sensitivity. Human Physiology, 34(2) 221–225. 10.1134/S0362119708020138

59. Xu, L., Han, Y., Chen, X. et al. (2020) Molecular mechanisms underlying menthol binding and activation of TRPM8 ion channel. Nat Commun 11, 3790. https://doi.org/10.1038/s41467-020-17582-x

60. Modern Ophthalmology (2009) A Guide. 2nd ed. Ed. V.F. Danilichev. SPb. Piter.

61. Filingeri, D., Havenith, G. (2015) Human skin wetness perception: psychophysical and neurophysiological bases, Temperature, 2(1) 86–104, https://doi.org/10.1080/23328940.2015.1008878

62. Bagchi, B. (2013) Water in Biological and Chemical Processes: From Structure and Dynamics to Function.

63. Matveev, V.V. (2019) Cell theory, intrinsically disordered proteins, and the physics of the origin of life, Prog. Biophys. Mol. Biol. 149, 114–130, https://doi.org/10.1016/j.pbiomolbio.2019.04.001

64. Kholmanskiy, A. (2020) Supramolecular physics of liquid water, Trends Phys. Chem. 20. 81–96, http://www.researchtrends.net/tia/abstract.asp?in=0&vn=20&tid=16&aid=6636&pub=2020&type=3

65. Kholmanskiy, A. (2021) Synergism of dynamics of tetrahedral hydrogen bonds of liquid water, Phys. Fluids. 33, 067120; https://doi.org/10.1063/5.0052566

66. Bardik, V., et al., (2020) The crucial role of water in the formation of the physiological temperature range for warm-blooded organisms, J. Mol. Liq. 306 112818, https://doi.org/10.1016/j.molliq.2020.112818.

